# High-Field Multinuclear MRI Reveals Sodium Relaxation Heterogeneity in Cortical Organoids

**DOI:** 10.64898/2026.04.01.715894

**Authors:** Grace Yu, Xiaochen Liu, David Hike, Chunqi Qian, Anna Devor, Ella Zeldich, Martin Thunemann, Xiaoqing Alice Zhou

**Affiliations:** Brain Rejuvenation and Vitality Enhancement (BRAVE) Laboratory, Athinoula A. Martinos Center for Biomedical Imaging, Massachusetts General Hospital and Harvard Medical School, Charlestown, MA, 02129; Translational Neuroimaging and Neural Control Laboratory, Athinoula A. Martinos Center for Biomedical Imaging, Massachusetts General Hospital and Harvard Medical School, Charlestown, MA, 02129; Department of Radiology, Michigan State University, East Lansing, MI, USA; Department of Electrical and Computer Engineering, Michigan State University, East Lansing, MI, USA; Department of Biomedical Engineering, Boston University, Boston, MA, USA; Department of Anatomy and Neurobiology, Boston University School of Medicine, Boston, MA, USA

## Abstract

Sodium magnetic resonance imaging (^23^Na MRI) provides a unique opportunity to probe ionic microenvironments in neural tissue because sodium ions play central roles in membrane electrophysiology, ion transport, and cellular homeostasis. Unlike conventional proton (¹H) MRI, which primarily reflects water distribution and tissue structure, ²³Na MRI is sensitive to ionic compartmentation and quadrupolar interactions arising from the spin-3/2 nature of the sodium nucleus. However, sodium MRI remains technically challenging due to intrinsically low signal sensitivity and rapid biexponential relaxation, particularly when imaging small biological systems. Here, we establish a high-field multinuclear MRI platform for imaging human cerebral organoids at 14 Tesla. Cerebral organoids derived from human induced pluripotent stem cells provide a simplified three-dimensional neural tissue model that enables investigation of ionic microenvironments without vascular or systemic confounds. Using a dual-tuned ¹H/²³Na radiofrequency coil, we performed co-registered structural, diffusion, and sodium imaging of individual fixed organoids. High-resolution ¹H MRI (33–100 μm) revealed pronounced microstructural heterogeneity, while multi-echo ²³Na MRI (300–400 μm) enabled voxel-wise characterization of quadrupolar relaxation behavior. Bi-exponential analysis of the sodium signal decay identified distinct relaxation components (T2*_short_ ≈ 1 ms and T2*_long_ ≈ 12 ms) and revealed spatial heterogeneity in sodium microenvironments across the organoid tissue. These results demonstrate the feasibility of quantitative sodium relaxometry in cortical organoids and establish a multinuclear imaging platform for investigating ionic microenvironment dynamics in three-dimensional neural tissue models.

## Introduction

Human organoids derived from pluripotent stem cells have emerged as powerful three-dimensional model systems for investigating disease mechanisms, drug screening, and therapeutic responses *in vitro*^1–3^. Cortical organoids recapitulate key aspects of neuronal differentiation, cellular diversity, and tissue organization, making them valuable platforms for studying neurodevelopmental and neurodegenerative disorders^4, 5^. Non-destructive imaging plays a central role in characterizing these systems across spatial scales while preserving their suitability for longitudinal experiments. Among available modalities, magnetic resonance imaging (MRI) offers a unique combination of deep tissue penetration, label-free contrast, and quantitative sensitivity to tissue structure, diffusion, and biomechanics. Recent studies have demonstrated that ultra-high-field MRI (9.4–28.2 T) can resolve organoid microstructure with spatial resolutions approaching tens of micrometers using diffusion MRI and T₂-weighted imaging^6–8^. These approaches enable visualization of internal architectural features such as rosette structures, cyst formation, and anisotropic tissue organization, and have been applied to monitor organoid growth, characterize disease phenotypes, and assess biomechanical properties^9, 10^. Collectively, these studies establish MRI as a promising platform for non-destructive organoid characterization and longitudinal monitoring.

In parallel with MRI-based studies, optical microscopy has become the dominant approach for interrogating cellular organization and functional activity in organoids. Confocal and multiphoton microscopy enable visualization of organoid cytoarchitecture, cellular differentiation, and protein localization through fluorescent labeling of neuronal and glial markers, providing detailed maps of cell-type distribution and tissue organization within three-dimensional organoid systems^3, 11–14^. These platforms can also be integrated with automated high-content imaging workflows capable of analyzing hundreds to thousands of organoids in multi-well formats, enabling scalable drug-screening and phenotypic profiling applications^15–18^. Furthermore, fluorescence-based calcium imaging has revealed spontaneous synchronized neuronal activity in midbrain or cortical organoids^15, 19^ and developmental maturation of intracellular Ca²⁺ signaling dynamics in kidney and ureteric organoids^20, 21^. While these fluorescence-based approaches provide powerful tools for monitoring neuronal activity and intracellular signaling pathways, they rely on reporter-based measures that primarily probe specific molecular pathways.

Complementary strategies for biochemical and metabolic characterization of organoids have therefore been explored using nuclear magnetic resonance (NMR) spectroscopy and related label-free imaging techniques. Hyperpolarized ¹³C NMR enables real-time monitoring of metabolic activity such as lactate dehydrogenase flux in cortical organoids^22^, while high-resolution magic-angle spinning (HR-MAS) spectroscopy allows non-destructive quantification of metabolites in intact organoids^23^. Spatiotemporal NMR spectroscopy has further revealed metabolic gradients within three-dimensional spheroids, including elevated lactate and reduced glucose concentrations in hypoxic cores^24^. Despite these advances in metabolic imaging, direct mapping of ionic microenvironments in intact organoids remains largely unexplored, and applications of ²³Na MRI to organoid systems have not yet been reported.

Among X-nuclei accessible by MRI, ²³Na offers a particularly compelling probe of ionic homeostasis, because sodium ions play fundamental roles in membrane excitability, ion transport, and metabolic regulation^25–27^. The quadrupolar relaxation properties of the spin-3/2 sodium nucleus further provide sensitivity to molecular ordering and microstructural environments that are not accessible through conventional proton MRI^28, 29^. Here we establish a multinuclear MRI platform for cortical organoid imaging that integrates high-resolution ¹H structural and diffusion MRI with quantitative ²³Na MRI using a custom dual-tuned RF coil at 14 T. This approach enables co-registered mapping of organoid microstructure, water diffusion properties, and sodium quadrupolar relaxation dynamics within the same sample. We demonstrate that diffusion MRI reveals heterogeneous tissue organization across fixed organoids, while multi-echo ²³Na imaging enables voxel-wise estimation of T2*_short_ and T2*_long_ relaxation components associated with distinct ionic microenvironments. Together, these results establish cortical organoids as a controllable experimental platform for multinuclear MRI investigations and provide a foundation for future studies of ionic microenvironment dynamics and sodium-based functional MRI mechanisms^30^.

## Methods

### Cortical organoid preparation

Human cortical organoids were generated from human induced pluripotent stem cells (hiPSCs) using an established differentiation protocol, following procedures described in our previous work^19, 31^. hiPSCs (WC-24-02-DS-M) were maintained under feeder-free conditions and dissociated to form embryoid bodies in ultra-low-attachment 96-well culture plates. On day 16, the organoids were transferred to the static 24-well ultra-low-attachment plates and cultured individually until harvested and fixed in 4% paraformaldehyde (PFA) for the experiments. From day 7 to 25, the media was supplemented with fibroblast growth factor 2 (FGF2; 20 ng/mL, cat. 233-FB-25/CF, R&D Systems, Minneapolis, MN, USA) and epidermal growth factor (EGF; 20 ng/mL, cat. 236-EG-200, R&D Systems) and with brain-derived neurotrophic factor (BDNF; 20 ng/mL, cat. AF-450-02, PeproTech, Waltham, MA, USA), neurotrophic factor 3 (NT3; 20 ng/mL, cat. 450-03, PeproTech) between days 25 to 39. No growth factors were added after day 40.

For MRI experiments, cortical organoids were collected at 12–13 weeks of differentiation, when the organoids typically reached ∼1.5–2 mm in diameter, providing sufficient tissue volume for multinuclear MRI measurements while preserving structural integrity. The fixed organoids were then embedded in 1–1.5% low-melting-temperature agarose (V2111, Promega) inside 5-mm NMR tubes filled with phosphate-buffered saline (PBS), which provided mechanical stabilization during MRI acquisition while maintaining hydration of the samples.

All experiments involving human induced pluripotent stem cells and derived cortical organoids were conducted in accordance with institutional biosafety and ethical guidelines and were approved by the Institutional Biosafety Committees (IBC) at Boston University and Massachusetts General Hospital.

### RF Coil Design

Because the intrinsic sensitivity of ²³Na MRI is substantially lower than that of ¹H MRI, maximizing radiofrequency (RF) detection efficiency was essential. We therefore fabricated a miniature solenoid RF coil (5 mm inner diameter) optimized for imaging small cortical organoids. The solenoid geometry provides high filling factor and improved sensitivity for small samples relative to conventional loop coils. The RF coil was implemented as a compact RLC resonant circuit composed of the solenoid inductor and a combination of fixed and variable capacitors to achieve resonance tuning and impedance matching to 50 Ω (Fig 1A)^32, 33^. The coil was tuned to the ²³Na Larmor frequency (∼158 MHz at 14 T) and ¹H frequency (∼599 MHz). To enable multinuclear imaging, a dual-tuned ¹H/²³Na configuration was constructed by integrating two resonance pathways within the same coil structure, allowing acquisition of both nuclei from the same sample without repositioning (Fig. 1B). To minimize electromagnetic coupling between the two RF channels, additional band-pass filtering and isolation were implemented along the RF transmission lines. Coil resonance characteristics and impedance matching were verified under in situ operating conditions using the Bruker system, with representative reflection (“wobbling”) curves shown in Fig. 1C. The RF coil was interfaced with a Bruker AVANCE Neo console, enabling independent RF transmission and reception for both the ¹H and ²³Na channels.

**Figure 1.**
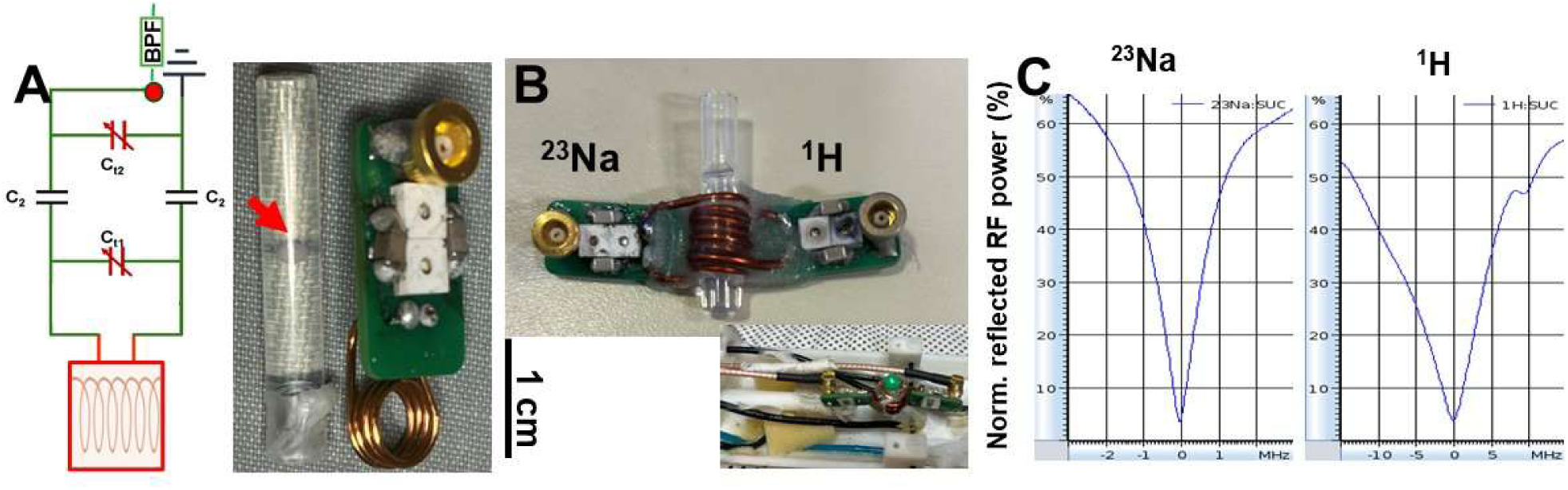
**Design and characterization of the dual-tuned ¹H/²³Na RF coil for organoid MRI.** A. Circuit schematic and photograph of the miniature solenoid RF coil used for imaging cerebral organoids. The coil forms an RLC resonant circuit consisting of the solenoid inductor and tuning/matching capacitors for impedance matching. B. Photograph of the dual-tuned RF coil integrated with a 5 mm NMR tube containing the organoid sample, enabling multinuclear imaging of ²³Na and ¹H signals without repositioning. An additional view shows the coil mounted within the MRI probe setup. C. Representative reflection (“wobbling”) curves showing the resonance characteristics of the RF coil tuned to the ²³Na (∼158 MHz) and ¹H (∼599 MHz) frequencies, confirming proper tuning and impedance matching for multinuclear MRI acquisition.

### MRI Acquisition

MRI systems: All experiments were performed on horizontal bore MRI scanners (Magnex Scientific, UK, 130 mm bore size) at the Athinoula A. Martinos Center for Biomedical Imaging (Charlestown, MA, USA). The 14 Tesla (∼600 MHz for ¹H, ∼158 MHz for ^23^Na) system was equipped with a Bruker Avance NEO console running ParaVision 360 v3.3 and a micro-imaging gradient set (Resonance Research, Inc.) providing peak gradient strength 1.2 T/m over a 60 mm diameter.

For dual-tuned ^1^H/^23^Na MRI, 3D-RARE T₂-weighted images were applied with the following parameters: TR/TE, 1s/8.54 ms; RARE factor, 8; NA, 4 ; acquisition BW, 25kHz, FOV, 6.4 × 6.4 × 6.4 mm³, matrix size, 128 × 128 × 128, with 50µm isotropic resolution for a total of ∼4 h 33 min. The diffusion-weighted ¹H MRI was acquired using a Bruker DtiStandard spin-echo diffusion sequence with the following parameters: TR/TE, 0.5s/20ms; NA, 2. Acquisition BW, 50kHz; FOV, 6.4 × 6.4 × 6.4 mm³; matrix size, 40 × 40 × 40; with 160 µm isotropic resolution for a total of ∼7 h 7 min. Diffusion preparation was implemented in spin-echo mode with a monopolar diffusion gradient scheme, diffusion gradient duration δ = 4 ms, and gradient separation Δ = 10 ms, with b-values = 500, 1500, 3000, 4500, 6000, 7500, and 9000 s/mm² for the ADC mapping. The sodium acquisition used the 3D gradient echo (GRE) ²³Na MRI sequence with the following parameters: TR/TE, 50 ms/3.2 ms; NA, 64; FA, 90°; FOV, 6.4 × 6.4 × 6.4 mm³; matrix size, 20 × 20 × 20, with 320 µm isotropic resolution for a total of ∼42 min 40 s. For T2* mapping, the 3D GRE ^23^Na MRI sequence with different TEs were used with the following parameters: TR/TEs, 100ms/0.68, 0.84, 1, 1.25, 1.5, 2, 3, 4, 5, 6.5, 8, 10, 15, 20, 30, 40, 50, 65, 80ms; NA, 24; FA, 90°; Acquisition BW, 20kHz; FOV, 6.4 × 6.4 × 6.4 mm³; matrix size, 16 × 16 × 16, with 400 µm isotropic resolution for a total of ∼20 min 29 s, respectively. For single solenoid RF coil (^1^H or ^23^Na), 3D-RARE T₂-weighted images were applied with the following parameters: TR/TE, 0.5s/12ms, rare factor, 8, matrix, 192 x 192 x 240, FOV, 6.4 x 6.4 x 8mm^3^, resolution, 33 µm³ isotropic, BW, 30kHz, NA, 9, total imaging time: 6 hours 36 mins.

And, diffusion-weighted imaging (DWI) sequence was used with the following parameters: TR/TE, 0.5s/20ms, matrix, 64×64×80, FOV, 6.4×6.4×8mm^3^, resolution, 100 µm³ isotropic, BW, 50kHz, b = 1,000, 2000, 3000, 4500, 6000, 7500, 10,000 s/mm², total imaging time: 4 hours 58 mins, for apparent diffusion coefficient (ADC) mapping. For ²³Na MRI T2* mapping, we applied two mapping strategies to measure the single quantum quadrupolar moment T_2_* decay: i). TR/TEs,100ms/0.68…80ms ; FA, 90°, NA, 24; acquisition BW, 25 KHz; matrix size, 27 x 20 x 20, FOV, 8.1 x 6 x 6mm^3^, with 300µm isotropic resolution for a total of 16 min, respectively; ii) TR/TEs,100ms/0.68…80ms ; FA, 90°, NA, 24; acquisition BW, 20 KHz; matrix size, 16 x 16 x 20, FOV, 6.4 x 6.4 x 8mm^3^, with 400µm isotropic resolution for a total of 12 min 48 s, respectively.

### Data Processing and Statistical Analysis

Image processing and quantitative analysis were performed using AFNI (Analysis of Functional NeuroImages) and MATLAB (MathWorks). AFNI was used for image visualization, alignment, and region-of-interest (ROI) definition, while MATLAB was used for model fitting and quantitative parameter estimation.

Diffusion MRI Analysis: *Diffusion-weighted images were processed to generate apparent diffusion coefficient (ADC) maps. Signal intensities acquired at different diffusion weightings were fitted voxel-wise using a monoexponential decay model:*

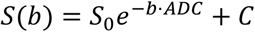

*where* 𝑆(𝑏)*represents the signal intensity at diffusion weighting* 𝑏, 𝑆_O_*denotes the signal without diffusion weighting (*𝑏 = 0*), and ADC is the apparent diffusion coefficient. C accounts for a constant noise floor. Voxel-wise fitting was performed in MATLAB using nonlinear least-squares* optimization to estimate ADC values for each voxel.

Sodium MRI Relaxation Analysis: Voxel-wise ²³Na signal decay curves acquired at multiple TEs were analyzed to characterize the quadrupolar relaxation properties of sodium in different microenvironments. The transverse magnetization signal M_xy_(t)was fitted using a biexponential decay model:

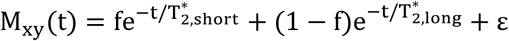

where frepresents the fractional contribution of the fast-decaying component, T^∗^ (typically 0.2–2 ms) corresponds to the rapidly decaying sodium component, and T^∗^(typically 5–20 ms) represents the slowly decaying component. εrepresents the residual fitting error. Parameter estimation was performed voxel-wise using nonlinear least-squares fitting implemented in MATLAB, generating quantitative T₂*_short_, T₂*_long_, and fractional component maps. The two relaxation components were interpreted as reflecting sodium ions residing in distinct microenvironments, such as restricted macromolecular-associated sodium (fast component) and relatively free sodium pools (slow component).

ROI Analysis: For quantitative analysis, regions of interest (ROIs) were manually defined on the anatomical images and applied to the corresponding ADC and sodium relaxation maps. Mean parameter values were extracted from each ROI for subsequent statistical analysis.

Statistical Analysis: All quantitative results are reported as mean ± standard error of the mean (SEM). For ROI-based comparisons of ADC values between conditions, paired two-tailed Student’s t-tests were performed. Statistical significance was defined as p < 0.02.

## Results

### Co-registered ¹H/²³Na MRI of Cortical Organoids Using Dual-Tuned RF Coils

Using the dual-tuned ¹H/²³Na RF coil, we performed multinuclear MRI of fixed cortical organoids at 14 Tesla, enabling co-registered T₂-weighted (50 μm isotropic) and diffusion-weighted (160 μm isotropic) ¹H MRI together with T₂-weighted ²³Na MRI (300 μm isotropic)* from the same sample (Fig. 2A). Despite differences in signal-to-noise ratio (SNR) between the two nuclei, the organoids were clearly delineated from the surrounding agarose gel based on their distinct relaxation properties. Specifically, organoids exhibited higher diffusion-weighted ¹H signals and lower T₂-weighted ²³Na signals* compared with the agarose gel, providing clear contrast between tissue and embedding medium (Fig. 2A). Diffusion-weighted images were acquired across multiple b-values to further characterize microstructural properties. As diffusion weighting increased, DWI signals progressively decreased, reflecting water diffusion within the organoid tissue (Fig. 2B). Fitting the signal decay with a mono-exponential diffusion model enabled estimation of apparent diffusion coefficient (ADC) maps, which revealed spatial heterogeneity across the organoids (Fig. 2C). In parallel, ²³Na MRI was acquired at multiple echo times (TE) to probe sodium relaxation properties within the organoids (Fig. 2D). The sodium signal exhibited rapid decay with increasing TE, consistent with the quadrupolar relaxation behavior of ²³Na nuclei. Together, these results establish a multinuclear MRI platform capable of co-registered structural, diffusion, and sodium imaging of cortical organoids.

**Figure 2.**
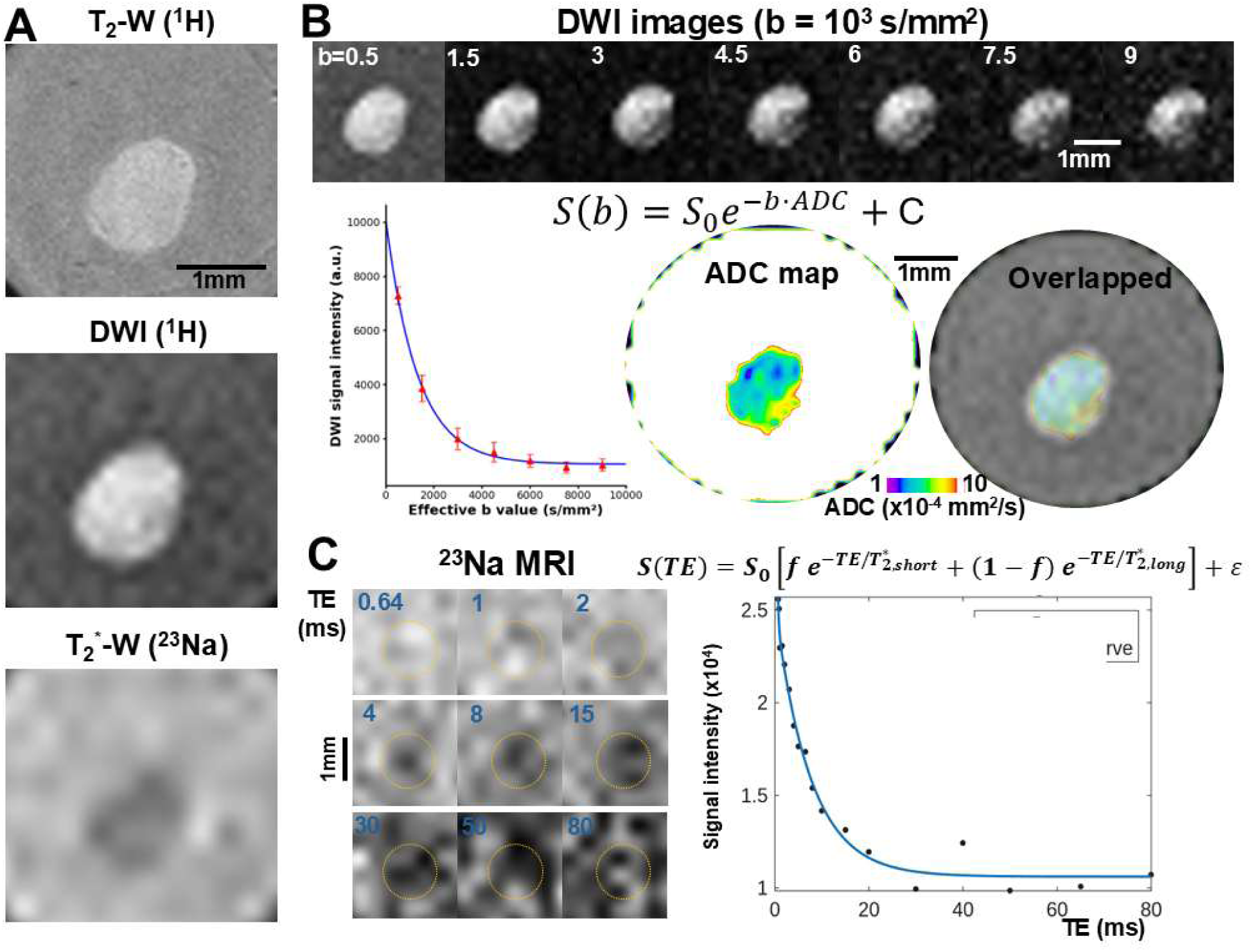
**Multinuclear MRI of cerebral organoids at 14 Tesla.** A. Co-registered multinuclear MRI of cerebral organoids acquired using a dual-tuned ¹H/²³Na RF coil. High-resolution ¹H T₂-weighted RARE images reveal the overall organoid morphology and structural organization. Diffusion-weighted imaging (DWI, b = 0.5 × 10³ s/mm²) provides diffusion-sensitive contrast highlighting differences in water mobility within the organoid tissue. ²³Na MRI acquired at short echo time (TE) provides sodium-weighted contrast from the same organoid preparation. B. Representative DWI images acquired at multiple b-values, showing progressive signal decay with increasing diffusion weighting. The signal attenuation reflects restricted water diffusion within the three-dimensional organoid tissue. Apparent diffusion coefficient (ADC) maps derived from mono-exponential fitting of the diffusion signal decay. Spatial variations in ADC values in color-coded maps reveal microstructural heterogeneity within the organoid. C. Multi-echo ²³Na MRI demonstrating sodium signal decay across increasing echo times. The rapid signal attenuation reflects the quadrupolar relaxation properties of the spin-3/2 sodium nucleus. Biexponential fitting enables characterization of distinct relaxation components associated with different sodium microenvironments.

### Diffusion MRI Reveals Microstructural Heterogeneity Within Organoids

To further characterize the internal microstructure of cortical organoids, we performed high-resolution ¹H MRI at 33 μm isotropic resolution (Fig. 3A). The T2-weighted RARE images revealed detailed structural contrasts within the organoids, highlighting heterogeneous internal features across different regions. DWI MRI further provided sensitivity to water diffusion through the tissue microstructure, displaying clear signal contrast reflecting differences in H2O diffusion across distinct microstructural environments within the organoids (Fig. 3B). These diffusion properties were quantified by estimating apparent diffusion coefficient (ADC) maps, which revealed spatial variations in diffusion across different organoids (Fig. 3C). Region-of-interest (ROI) analyses demonstrated distinct b-value-dependent signal decay profiles, indicating differences in diffusion behavior between regions. Consistent with these observations, quantitative analysis showed significantly different ADC values across organoid regions (Fig. 3D), supporting the presence of microstructural heterogeneity within the organoids.

**Figure 3.**
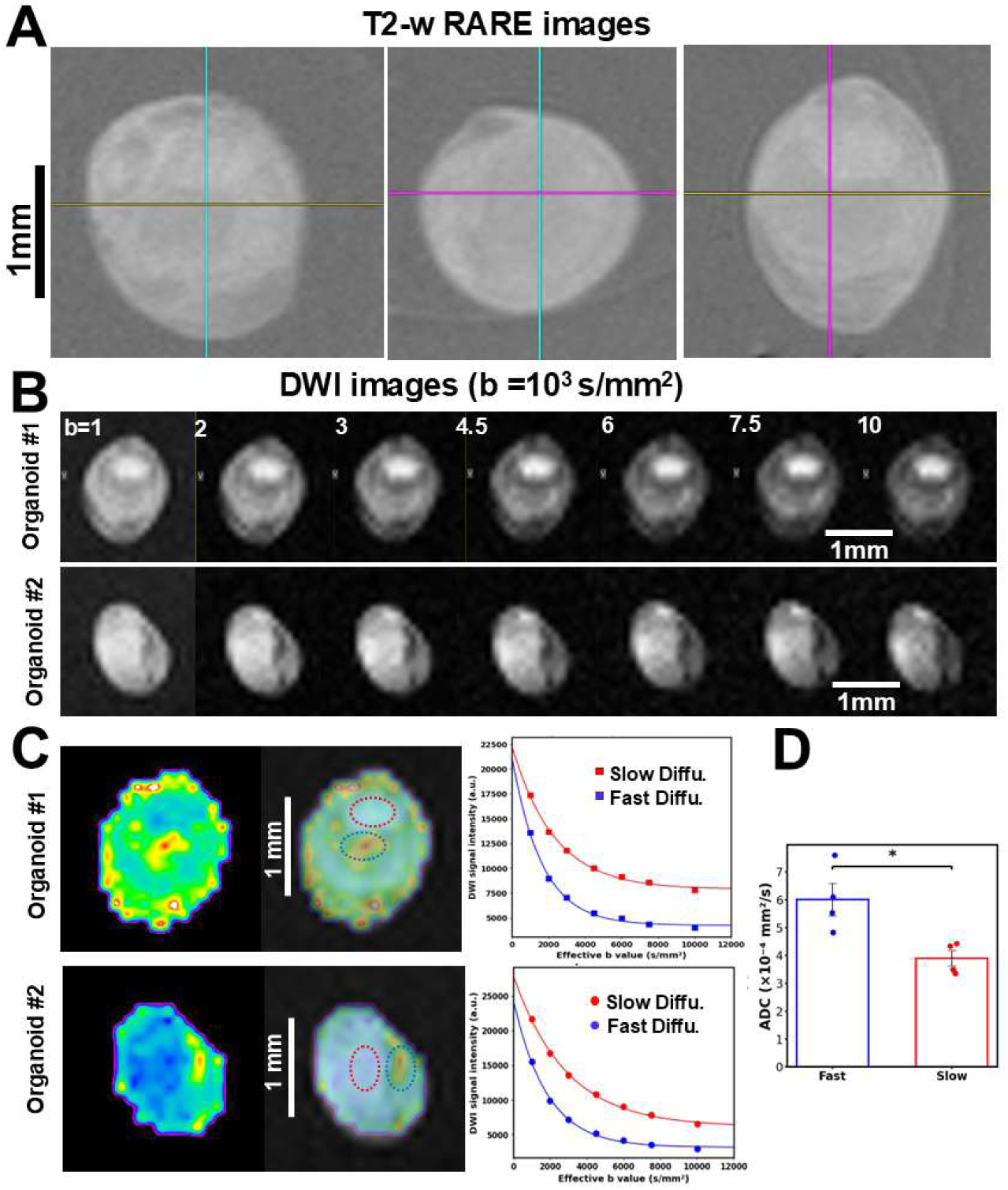
**High-resolution ¹H MRI reveals microstructural heterogeneity in cerebral organoids** A. High-resolution ¹H T₂-weighted RARE images of a cerebral organoid acquired at 33 μm isotropic resolution, shown in three orthogonal views. The images reveal detailed internal structural organization and heterogeneous signal contrast across the organoid. B. Diffusion-weighted images (DWI) acquired at increasing b-values from two representative organoids, showing progressive signal attenuation associated with water diffusion within the tissue. c, Apparent diffusion coefficient (ADC) maps derived from diffusion signal fitting. Spatial variations in ADC values highlight heterogeneous diffusion properties within the organoid. Representative regions of interest (ROIs, dashed circles) exhibit distinct diffusion signal decay profiles across b-values. d, Quantitative comparison of ADC values across ROIs, demonstrating significant diffusion heterogeneity within the organoid tissue (n = 4 organoids, Fast ADC: 6.02±0.59×10^−4^mm^2^/s; Slow ADC: 3.91±0.29×10^−4^mm^2^/s; *p < 0.02, paired two-tailed Student’s t-test). Error bars represent SEM.

### ²³Na MRI Reveals Quadrupolar Relaxation Heterogeneity in Cortical Organoids

To investigate ionic microenvironments within the organoids, we performed ²³Na MRI using the dual-tuned RF coil at 300–400 μm isotropic resolution. Representative *T₂-weighted ²³Na images (TE = 1 ms)* from three consecutive slices clearly delineated the organoid structure relative to the surrounding agarose (Fig. 4A). To further characterize sodium relaxation behavior, multi-echo ²³Na acquisitions were performed across a range of echo times (TE), enabling voxel-wise estimation of quadrupolar T2* decay. Using biexponential fitting, we derived spatial maps of T2*_short_ and T2*_long_ components across the organoids. The resulting color-coded relaxation maps revealed region-specific variations in sodium relaxation properties, reflecting heterogeneous ionic microenvironments associated with different tissue microstructures (Fig. 4A).

**Figure 4.**
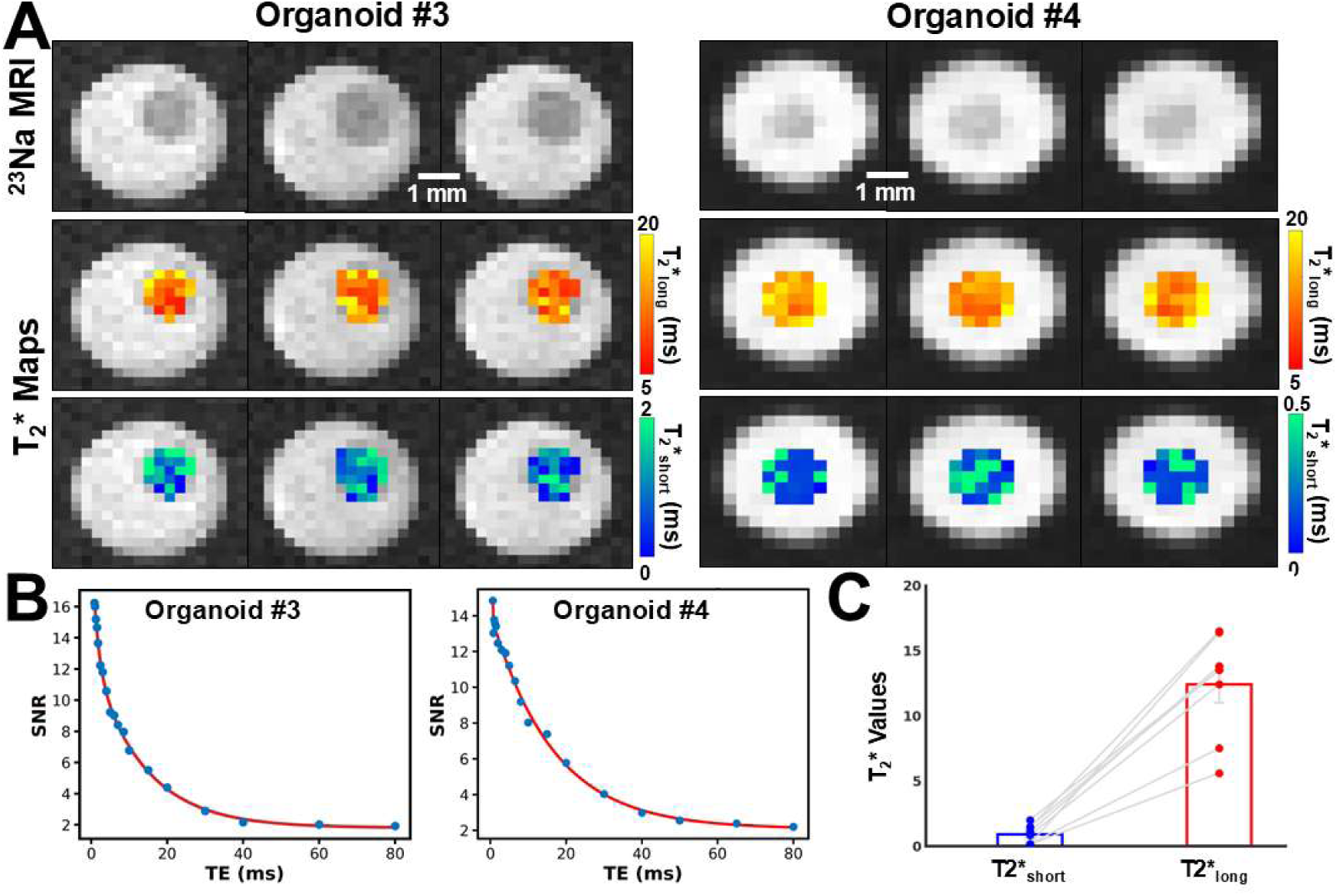
**²³Na MRI reveals biexponential relaxation and ionic microenvironment heterogeneity in cerebral organoids** A. Representative ²³Na MRI images from two cerebral organoids acquired at TE = 1 ms, shown across three consecutive slices. Corresponding color-coded T2\**_short_ and T2\**_long_ maps derived from multi-echo sodium imaging are overlaid on the anatomical sodium images. The maps illustrate spatial variations in sodium quadrupolar relaxation properties within the organoid tissue. B. Representative ²³Na quadrupolar decay curves from two organoids showing signal attenuation across increasing echo times. Biexponential fitting of the decay curves yields distinct relaxation components corresponding to fast and slow sodium environments. C. Group-level quantification of sodium relaxation components across organoids. Bar plots show mean T2*_short_ (0.91 ± 0.25 ms) and T2*_long_ (12.4 ± 1.38 ms)(mean ± SEM, n = 8 organoids), demonstrating reproducible biexponential sodium relaxation consistent with heterogeneous ionic microenvironments.

Representative quadrupolar decay curves and biexponential fits are shown in Fig. 4B, demonstrating the presence of fast and slow relaxation components consistent with the spin-3/2 quadrupolar nature of ²³Na nuclei. Group-level analysis of the estimated relaxation parameters showed reproducible T2*_short_ *and T2\**_long_ values across organoids (Fig. 4C), supporting the presence of heterogeneous sodium environments within the organoid tissue. Together, these results establish cortical organoids as a controllable platform for multinuclear MRI, enabling integrated ¹H structural imaging and ²³Na-based ionic mapping, with potential applications in X-nuclei MRI for pharmacological studies, drug screening, and investigation of sodium-based fMRI mechanisms.

## Discussion

This study establishes a multinuclear MRI platform for cortical organoids at ultra-high magnetic field, enabling co-registered structural, diffusion, and ionic imaging within the same tissue preparation. Using a custom dual-tuned ¹H/²³Na RF coil, we demonstrate that diffusion MRI can resolve microstructural heterogeneity within fixed individual organoids, while multi-echo ²³Na MRI enables quadrupolar relaxation mapping of ionic microenvironments through measurements of T2*_short_ and T2*_long_ components. Together, these results show that organoid MRI can provide complementary structural and ionic information in a controlled neural model system that lacks vascular and hemodynamic confounds. This capability creates new opportunities to study ionic regulation, neuronal activity–related sodium dynamics, and pharmacological perturbations using multinuclear MRI in living organoid systems.

### DWI-MRI reveals microstructural heterogeneity in cortical organoids

DWI-MRI provides a quantitative readout of tissue microstructure through measurements of the apparent diffusion coefficient (ADC), which reflects the mobility of water molecules within cellular environments. In the present study, ADC mapping revealed clear regional heterogeneity within cortical organoids, with region-specific values of approximately 0.39–0.60 × 10⁻³ mm²/s across different organoid regions (Fig 3). The ADC values measured here fall within the general range reported in recent organoid diffusion MRI studies, although they are slightly lower than those reported in a 28.2 T cortical organoid diffusion MRI study, which observed mean ADC values of approximately 0.81–0.97 × 10⁻³ mm²/s in unfixed organoids^7^. Differences in sample preparation likely contribute to this discrepancy. Diffusion measurements are highly sensitive to tissue state, including whether samples are living, unfixed ex vivo, or chemically stabilized.

It is also informative to compare these measurements with diffusion values reported in brain tissue. In vivo human brain ADC values are typically around 0.7–0.9 × 10⁻³ mm²/s in gray matter and 0.6–0.8 × 10⁻³ mm²/s in white matter at physiological temperature^34, 35^. By contrast, chemically fixed brain tissue exhibits substantially reduced diffusion coefficients, typically in the range of 0.2–0.5 × 10⁻³ mm²/s, reflecting reduced water mobility caused by tissue cross-linking and altered extracellular space^36, 37^. The ADC values measured in the present organoids fall between these regimes, suggesting that organoid tissue exhibits diffusion restriction intermediate between living brain tissue and fixed brain samples. The difference likely reflects variations in cellular packing density, extracellular space fraction, and local tissue organization during organoid development. Taken together, these comparisons indicate that diffusion MRI provides a sensitive marker of organoid microstructural organization, while the observed ADC values highlight fundamental differences between organoid cellular assembly and mature brain architecture. Importantly, the ability to detect spatially heterogeneous diffusion compartments within individual organoids provides a structural framework for interpreting the multinuclear MRI measurements of ionic microenvironments described below.

### 23Na quadrupolar mapping reveals ionic microenvironment heterogeneity in organoids

The second major advance of this work is the demonstration that ²³Na quadrupolar decay can be measured in cortical organoids with sufficient sensitivity to derive voxel-wise T2*_short_ and T2*_long_ maps. This is technically important because sodium MRI is intrinsically difficult: the observable ²³Na signal is far weaker than the proton signal because of both lower NMR sensitivity and much lower physiological concentration, while the spin-3/2 sodium nucleus undergoes rapid biexponential transverse relaxation due to quadrupolar interactions. As a result, robust estimation of T2*_short_ and T2*_long_ requires high SNR, very short echo-time sampling, and fitting strategies that remain stable under rapidly decaying signal conditions. In the classical fast-motion regime, the single-quantum sodium signal is often modeled as the sum of short and long components arising from the relative contributions of the central and satellite transitions, and many studies have used either a constrained biexponential model with fixed component fractions or a free-fraction biexponential fit depending on SNR and echo sampling density^38–41^. Early theoretical and experimental work by Rooney and Springer established the physical basis for interpreting ²³Na relaxation in terms of local molecular ordering, rotational restriction, and macromolecular interactions, while later quantitative MRI studies emphasized that biexponential relaxation must be explicitly considered to avoid bias in sodium concentration measurements^42–47^. Accurate estimation of T2*_short_ requires ultrashort echo-time sampling because the fast sodium component decays on the millisecond timescale. If early echoes are not captured, the biexponential fit becomes poorly constrained, particularly at low SNR. In conventional in vivo sodium MRI, this requirement often conflicts with spatial resolution because larger fields of view and lower sodium sensitivity limit achievable SNR at very short TE. In contrast, organoid imaging benefits from the small sample size and reduced field of view, which enables acquisition at relatively higher spatial resolution while maintaining adequate SNR and ultrashort TE sampling for robust estimation of T2*_short_ and T2*_long_. This combination makes organoid preparations particularly well suited for studying sodium quadrupolar relaxation behavior and for testing quantitative relaxometry models under controlled experimental conditions.

Our work provides T2* measurements of ^23^Na in organoids, presenting a critical comparison platform with the reported sodium relaxation values in the human brain. Across tissues, the short component is generally interpreted as reflecting sodium in more restricted, partially ordered, or protein-rich environments, whereas the long component reflects sodium in less restricted environments. Quantitative multi-echo sodium MRI studies of the human brain have reported T2*_short_ ≈ 3-4 ms and T2*_long_ ≈ 20-26 ms in white and gray matter using voxel-wise biexponential analysis, and have further shown that both components vary substantially across brain regions, consistent with sensitivity to tissue composition and compartmental organization^45, 48^. More broadly, sodium MRI studies summarize that biological tissues typically exhibit T2*_short_ in the range of ∼0.5-5 ms and T2*_long_ of ∼15-30 ms, depending on tissue type, field strength, and fitting model^43, 45, 49–52^. Against this background, the organoid measurements reported here, approximately ∼1 ms for T2*_short_ and ∼12-14 ms for T2*_long_, fall within the expected general range of biexponential sodium decay, with a short component near the lower end of the *in vivo* range and a shorter long component than typically observed in adult brain.

This pattern is biologically plausible and suggests that sodium in organoids experiences substantial microenvironmental restriction while lacking some of the weakly restricted long-T2* contributions present in mature brain tissue, where vascular, CSF, and larger extracellular compartments contribute to the overall signal.

### Potential applications of organoid multinuclear MRI

A primary motivation for establishing this platform is its potential relevance to the mechanistic interpretation of neuronal-activity-related sodium (NARS) fMRI^30^. A central question in sodium-based functional MRI is whether neuronal activation produces detectable sodium signal changes that reflect activity-dependent ionic dynamics rather than vascular or bulk concentration effects. *In vivo*, addressing this question is challenging because neuronal sodium fluctuations occur simultaneously with hemodynamic responses, blood volume changes, and cerebrospinal fluid partial-volume effects. Cortical organoids provide a simplified preparation in which neuronal tissue organization and ionic regulation can be studied without vascular and hemodynamic confounds. The present work establishes a key prerequisite for such investigations by demonstrating that sodium quadrupolar relaxometry can be measured robustly in organoid tissue, yielding quantifiable T2*_short_ and T2*_long_ components that report local ionic microenvironment. In this context, organoid ²³Na MRI provides a controlled experimental system for examining whether neuronal activity is accompanied by shifts in sodium relaxation behavior consistent with mechanisms proposed for NARS-fMRI.

The ability to perform multinuclear MRI in organoids may also enable exploratory studies of other X-nuclei relevant to cellular physiology, although such applications remain technically challenging. For example, ⁷Li MRI^53, 54^, which has been explored in the context of psychiatric treatment and intracellular signaling, represents one possible target for future investigation. While sensitivity limitations remain substantial, bipolar disease organoids could provide a compact and experimentally controllable system for evaluating acquisition strategies and quantitative models for nuclei with low intrinsic sensitivity^55, 56^.

More broadly, multinuclear organoid MRI may offer opportunities for studying ionic regulation and pharmacological perturbations in neural tissue models. Because ionic homeostasis and membrane transport are fundamental indicators of cellular viability, excitability, and metabolic state, MRI measurements of ion-sensitive contrast could potentially complement existing optical, electrophysiological, and molecular assays. In this framework, organoid MRI may serve as a non-destructive imaging approach for investigating drug responses, ion-channel function, and therapeutic modulation of neural tissue physiology within three-dimensional organoid systems.

## Conclusion

In this study, we developed and validated a 14 T dual-tuned ¹H/²³Na MRI platform for multinuclear imaging of human cortical organoids. High-resolution ¹H MRI provided structural and diffusion-based characterization of organoid microstructure, revealing spatial heterogeneity in tissue organization. In parallel, multi-echo ²³Na MRI enabled quantitative mapping of sodium quadrupolar relaxation, allowing estimation of T2*_short_ and T2*_long_ components that report ionic microenvironment heterogeneity within organoid tissue. Together, these results demonstrate the feasibility of performing quantitative multinuclear MRI in cortical organoids, integrating structural and ionic information within the same preparation.

Beyond establishing a technical platform, this work highlights the potential of organoid MRI as a controlled experimental system for studying sodium-based imaging mechanisms without vascular or hemodynamic confounds. By enabling quantitative characterization of sodium relaxation behavior in a simplified neural tissue model, the organoid ²³Na MRI framework provides a foundation for investigating activity-dependent ionic dynamics relevant to sodium-based functional MRI. More broadly, multinuclear MRI of organoids may support future studies of ionic regulation, pharmacological perturbations, and therapeutic responses in three-dimensional neural tissue models.

## Acknowledgements

This work was supported by the Alzheimer’s Association (ARFD-23-1145375 to X.A.Z.) and the Brain & Behavior Research Foundation Young Investigator Award (to X.A.Z.). Additional support was provided by NIH grants (RF1NS124778, R01NS122904, U19NS123717), NSF grant 2123971, and the S10 instrumentation grants (S10MH124733, S10OD036211) to the Martinos Center as well as NIH/NIA (RF1AG088529, awarded to E. Zeldich and M. Thunemann). We are also grateful to Dr. Jiang for the support in sharing the ²³Na MRI pulse sequences.

## Author Contributions

G.Y. performed experiments, analyzed data, and wrote the paper; X.L. analyzed the data, D.H. provided MRI technical support, C.Q. supervised RF coil design; A.D. built the concept; E. Z. provided the organoids, M.T. prepared organoids and acquired; X.A.Z. built the concept, supervised experiments and wrote the paper.

## Data and Code availability

The imaging datasets generated in this study will be deposited in a public repository prior to publication and made accessible via a unique accession code. Custom scripts with representative datassett used for preprocessing and bi-exponential modeling of T2* quadrupolar decay are deposited in a public repository (github: https://github.com/xl872/Na_T2.git).

